# Axon-specific mRNA translation shapes dopaminergic circuit development

**DOI:** 10.1101/2025.11.25.690389

**Authors:** C. Gora, B. Frenette, P. Gelon, C. F. Sephton, S.M.I. Hussein, E. Metzakopian, M. Lévesque

## Abstract

The precise organization of midbrain dopaminergic (mDA) projections is essential for motor and cognitive functions, and their disruption contributes to multiple brain disorders. Yet the molecular mechanism guiding the development of these projections remains poorly defined. Here, we used ribosome tagging (RiboTag) and specific mouse crossing strategies (DATIRES-Cre mice) to isolate ribosome-bound mRNAs specifically from mDA axons and to analyze their axonal translatomes across developmental stages. We found that early-stage axons are enriched in transcripts involved in axon guidance and growth, while mature axons predominantly translate mRNAs related to synaptic function. Among key candidates, we identified PlxnA4, which is locally translated into mDA axons and modulates arborization in response to Sema3a. Functional assays *in vitro* and *in vivo* revealed that Plxna4-mediated signaling regulates topographical axon targeting and innervation, particularly in the nigrostriatal pathway. Our results uncover a dynamic and compartment-specific regulation of mRNA translation in developing mDA neurons, offering mechanistic insight into circuit formation and providing new molecular targets to improve integration of grafted neurons in regenerative therapies for Parkinson’s disease.

## INTRODUCTION

The midbrain dopaminergic (mDA) system is composed of anatomically and molecularly distinct neuronal populations that project in a highly organized manner to specific forebrain targets (Poulin et al., 2014). This precise wiring underlies essential functions such as movement, reward processing, and executive control. Aberrations in dopaminergic (DA) circuitry are implicated in a broad spectrum of neurological and psychiatric conditions, notably Parkinson’s disease, schizophrenia, attention deficit disorders, autism, major depression, and drug addiction (Dickson, 2018; Huang et al., 2015).

Midbrain DA neurons are traditionally grouped into three anatomical clusters: the substantia nigra pars compacta (SNpc) (A9), the ventral tegmental area (VTA) (A10), and the retrorubral area (RR) (A8) (Björklund & Dunnett, 2007). These neurons give rise to distinct projection pathways, such as the nigrostriatal, mesolimbic, and mesocortical circuits, which respectively innervate the dorsal striatum, nucleus accumbens, and prefrontal cortex. Recent transcriptomic studies have revealed that mDA neurons are far more heterogeneous than previously appreciated, with molecular subtypes differing in gene expression, connectivity, and functional properties (Azcorra et al., 2023; Gaertner et al., 2022; Poulin et al., 2014). This diversity is thought to underline the broad range of DA functions and may contribute to the selective vulnerability observed in neurodegenerative diseases (Brichta & Greengard, 2014; Giguere et al., 2018; Lerner et al., 2015; Poulin et al., 2018; Surmeier et al., 2017). Despite their heterogeneity, mDA axons exhibit remarkable topographic organization, with specific subpopulations projecting to defined forebrain targets (Aransay et al., 2015; Chabrat et al., 2017; Prensa & Parent, 2001).

Although considerable progress has been made in mapping mDA circuitry, the molecular mechanisms guiding the development of these projections remain poorly understood. Both *in vitro* and *in vivo* studies in different central nervous system regions have shown that axons harbor distinct pools of mRNA that support axon growth and guidance (Douglas S. Campbell, 2001; Jung et al., 2023; Poulopoulos et al., 2019; Wu et al., 2005). In retinal ganglion cells, for instance, an axon-specific translatome has been described to support stage-specific processes such as elongation, branching, pruning, and synaptogenesis (Shigeoka et al., 2016). Axons also exhibit local translation of ribosomal proteins, which are incorporated into existing ribosomes, sustaining active protein synthesis far from the soma (Shigeoka et al., 2019). Additionally, RNA localization is tightly regulated by 3’ untranslated regions (3’ UTRs) and their interactions with RNA-binding proteins, which modulate mRNA distribution and remodeling during neuronal development (Luisier et al., 2023). Given the conservation of these mechanisms across species and neuronal types, we hypothesize that local translation plays a critical role in mDA axon development.

To explore this, we used the RiboTag mouse model (Sanz et al., 2009), in which ribosomes are tagged with hemagglutinin (HA) under the Cre-recombinase control. By crossing RiboTag mice with DAT-IRES-Cre mice, we selectively labeled ribosomes in dopamine transporter (DAT)-expressing neurons, enabling the isolation of ribosome-bound mRNAs from both mDA cell bodies and axons. Our analyses reveal axon-specific translatomes vary by projection target and developmental stage, suggesting that localized translation contributes to the spatial and temporal organization of mDA connectivity. These findings offer novel molecular insights into how dopaminergic circuits are assembled and may inform strategies to enhance circuit reconstruction in disease models.

## RESULTS

### RiboTag selectively labels ribosomes in dopaminergic neurons and their axons *in vivo*

To investigate the presence and identity of the mRNA within developing DA axons, we used the RiboTag mouse line (Sanz et al., 2009) in which Cre-mediated recombination replaces exon 4 of the ribosomal protein RPL22 with a hemagglutinin (HA)-tagged version (RPL22-HA). To restrict labelling to DA neuron, we crossed RiboTag mice with the DAT-IRES-Cre mice, which express Cre recombinase in DA neurons beginning at embryonic day 12. The resulting double transgenic animal, referred as DA-RiboTag mice, enables the isolation of ribosomes-bound mRNAs specifically from DA neurons and axons (**Fig. 1A**). We validated the specificity of HA-tagged ribosome expression in DA neurons and their axons using three complementary approaches. Immunohistochemistry on coronal sections of adult DA-RiboTag midbrain revealed HA immunoreactivity restricted to tyrosine hydroxylase (TH)-positive DA neurons (**Fig. 1B**). Importantly, no HA-positive cell bodies were detected in the striatum, confirming the absence of local HA expression by resident cells and supporting the specificity of mRNA labeling in dopaminergic projections. Because axonal HA levels are too low for conventional immunodetection (Hobson et al., 2022), we used proximity ligation assay (PLA) to detect HA-tagged ribosomes in DA axons with enhanced sensitivity. PLA detects proteins that are in proximity (<40 nm), producing a punctate fluorescent signal only when both target epitopes are co-localized. We performed PLA on E14.5 midbrain explants from DA-RiboTag and wild-type mice, using antibodies against RPL-P0, a core component of the large ribosomal subunit, and HA, which tags the genetically modified RPL22 in DA neurons. Punctate PLA signals were observed selectively within TH-positive axons in DA-RiboTag explants, but not in wild-type controls (**Fig. 1C**). To provide further molecular support for ribosome transport into axons, we conducted polyribosome fractionation from the midbrain and striatum of postnatal day 1 (P1) DA-RiboTag and wild-type mice. HA-tagged ribosomes were detected in both midbrain and striatal fractions from DA-RiboTag mice but were absent in wild-type controls (**Fig. 1D**), providing molecular evidence that ribosomes are actively transported into DA axons *in vivo*. Together, these histological, molecular and biochemical approaches indicated that the DA-RiboTag model enables selective labelling of ribosomes in both DA soma and their axonal projections, providing a robust tool to investigate axon-specific mRNA translation during dopaminergic circuit development.

**Figure 1.**
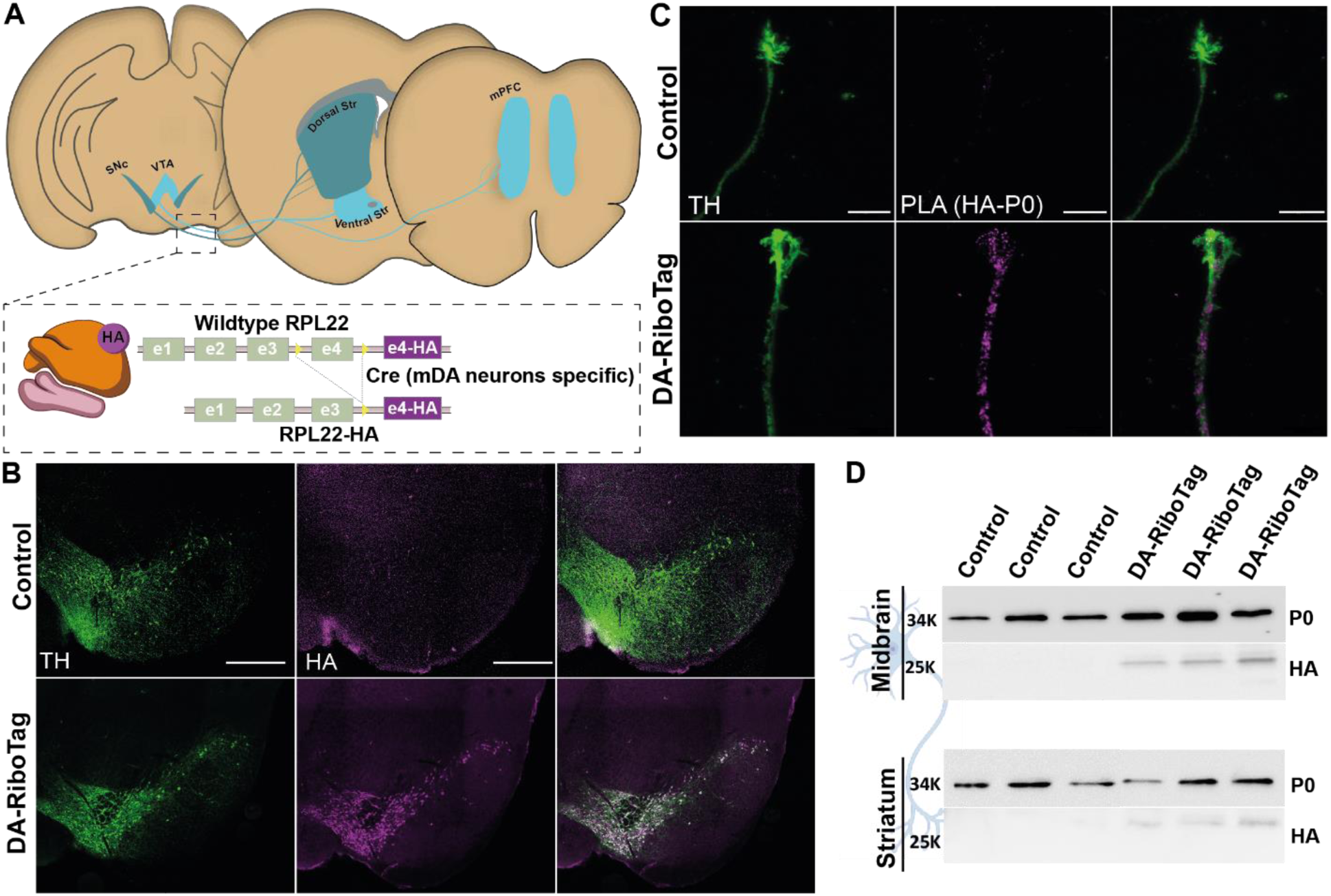
Successful labeling of dopaminergic axonal ribosomes in RiboTag mice crossed with Dat-IRES-Cre mice. A) Schematic representation of the dopaminergic projections and the axon-TRAP strategy specific to dopaminergic neurons. B) HA labeling of ribosomes in TH^+^ neurons (green) was revealed by HA immunohistochemistry (magenta). C) Representative images of proximity-ligation assay against HA and P0 in DA axons of midbrain explants from control and DA-Ribotag mice. D) Western-blot of one-day postnatal mouse midbrain and striatum, to reveal the expression of HA tag in Dat-IRES-Cre animals.

### RiboTag profiling reveals dynamic and projection-specific changes in ribosome-bound mRNAs across dopaminergic development

To understand how local translation shapes the development of DA projections, we profiled ribosomes-bound mRNAs in DA cell bodies and axons at distinct developmental stages. While previous studies have highlighted the molecular and anatomical heterogeneity of mDA neurons (Garritsen et al., 2023; Hook et al., 2018; La Manno et al., 2016; Poulin et al., 2020; Tiklova et al., 2019), less is known about how this diversity is reflected in the axonal translatome across space and time. Using the DA-RiboTag mouse model, we isolated HA-tagged ribosome-associated mRNAs from mDA somata in the midbrain and from axonal projections to the dorsal striatum, ventral striatum, and prefrontal cortex. Samples were collected at two time-points: P1, corresponding to the period of axon growth and branching, and adulthood (P60), when DA circuits have reached maturity. To selectively isolate DA axonal projections, we performed microdissections on fresh brain tissue, followed by mRNA extraction using the optimized Axon-TRAP protocol (Shigeoka et al., 2016). Because the HA-tagged ribosomal protein RPL22 is expressed only in DAT-expressing neurons in the DA-RiboTag model, only ribosome-bound mRNAs originating from DA neurons were captured.

We first compared the ribosome-bound mRNA profiles between P1 and adult samples using differential expression analysis to identify differentially expressed mRNAs (DE mRNAs) across each region. This analysis revealed that the number of DE mRNAs was consistently higher in axonal samples at P1 than in adulthood across all projection areas (**Fig. 2A-D**). In contrast, the mDA cell bodies in the midbrain exhibited relatively stable ribosome-associated mRNA profiles between developmental stages, suggesting that dynamic regulation of translation is more prominent in axons than in somata during development.

**Figure 2.**
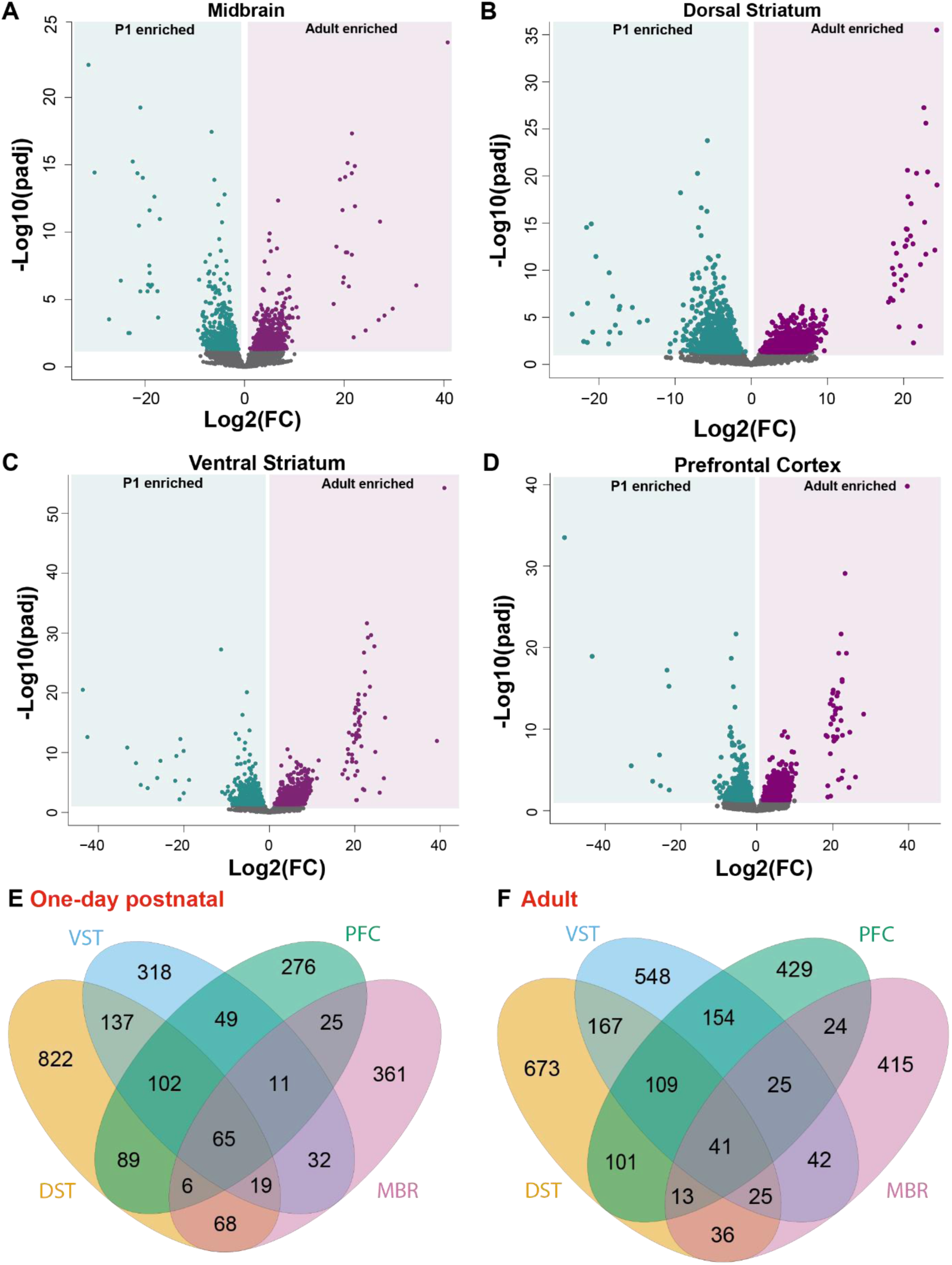
Identification of dopaminergic cell bodies and axonal translatome. A-D) Volcano plot of differentially expressed genes between two developmental stages: one-day postnatal and adult. Volcano plots are represented for each structure: midbrain, containing the DA cell bodies; dorso-ventral striatum and prefrontal cortex, containing DA axons. Blue dots are genes that are enriched at P1, and magenta dots are genes that are enriched in adult. Grey dots, genes that are not significantly enriched either in P1 or adult. E) Venn Diagram of genes that are significantly enriched in P1 from midbrain, dorsal striatum, ventral striatum and prefrontal cortex. F) Venn Diagram of genes that are significantly enriched in adult from midbrain, dorsal striatum, ventral striatum and prefrontal cortex.

Next, we examined how many of these DE mRNAs were unique to a given projection area or shared across structures. At P1, among the 2,380 DE mRNAs identified across all samples, 34.5% (822) were specific to mRNAs isolated from axons innervating the dorsal striatum, 13.4% (318) from axons targeting the ventral striatum, 11.6% (276) from axons projecting to the prefrontal cortex, and 15.2% (361) from DA cell bodies in the midbrain (**Fig. 2E**). A subset of DE mRNAs (16.8%, 400) was shared between two regions, 5.8% (138) were present in at least three compartments, and only 2.7% (65) were detected across all four sampled regions. These results underscore the spatial compartmentalization of mRNA translation in developing mDA neurons, with distinct axonal signatures depending on their projection targets. Notably, the somatic translatome remained largely distinct from that of axons, supporting the idea that local translation supports projection-specific developmental programs.

We then asked whether similar organizational principles applied to the mature DA system. At the adult stage, a comparable pattern emerged: among the 2,802 DE mRNAs identified, 24% (673) were specific to mRNAs isolated from axons projecting to the dorsal striatum, 19.6% (548) from axons innervating the ventral striatum, 15.3% (429) from axons targeting the prefrontal cortex, and 15.2% (415) from DA cell bodies in the midbrain (**Fig. 2F**). While some overlap in DE mRNAs were observed across compartments, the mature mDA system maintained a high degree of target-specific and spatially compartmentalized mRNA translation. Together, these results suggest that DA axons exhibit developmentally regulated patterns of ribosome-bound mRNAs that vary according to their projection targets and differ from those found in somata. These findings are consistent with the idea that axon-specific translation contributes to circuit formation during development and may be sustained in the adult brain to support projection-specific functions.

### DA axonal translatome shifts from axon elongation to neurotransmission during development

To determine which classes of mRNAs were preferentially associated with ribosomes in DA cell bodies and axons at different developmental stages (P1 or adult), we performed a gene ontology (GO) enrichment analysis on the DE mRNAs identified for each DA projection area and in the midbrain. Analyses were conducted separately for each compartment and developmental time points. GO terms related to cellular components showed considerable overlap between axonal and somatic compartments (**Fig. S1**), consistent with the idea that many mRNAs produced in the DA soma are trafficked into axons. Importantly, axonal samples also showed enrichment for cellular component terms related to axon structure and function, supporting the presence of compartment-specific translation processes in distal axons. In contrast, biological process GO terms revealed a clear shift in the axonal translatome across development. At P1, axons targeting all forebrain regions, as well as DA cell bodies, were enriched for terms associated with axon development, including axonogenesis, cytoskeletal dynamics, and guidance-related processes (**Fig. 3 A-D & Fig. S2-S3**). In adult samples, the predominant enrichment shifted toward synaptic functions such as synapse organization, neurotransmitter release, and regulation of synaptic plasticity (**Fig. 3 A-D**). These findings are consistent with a developmental transition in local mRNA translation, from supporting axon outgrowth to modulating mature synaptic function.

**Figure 3.**
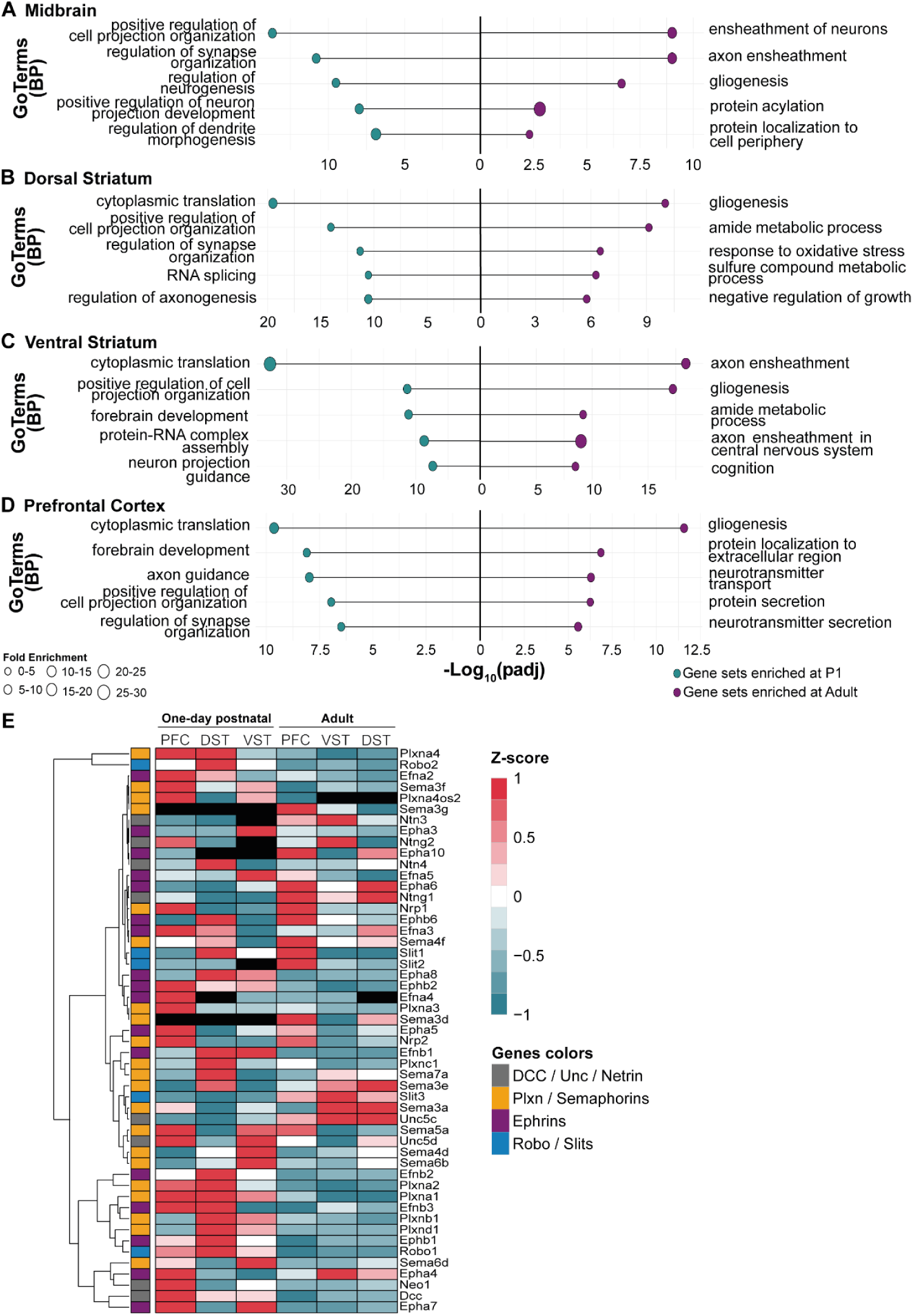
Evolution of biological process from young to mature dopaminergic neurons and axons. Go-term analysis showed the 5 representatives Go-term identify in each dopaminergic compartment. A) Go-term analysis of mRNAs presents in DA cell bodies at one-day postnatal and adult. B) Go-term analysis of mRNAs presents in DA axon projections to the dorsal striatum at one-day postnatal and adult. C) Go-term analysis of mRNAs presents in DA axon projections to the ventral striatum at one-day postnatal and adult. D) Go-term analysis of mRNAs presents in DA axon projections to the prefrontal cortex at one-day postnatal and adult. E) Z-score of mRNA expressions within DA axons in developmental stages P1 and adult. Data are represented in z-score of the mean of normalized count.

To explore whether axon guidance mechanisms are differentially regulated during development, we examined the expression of mRNAs encoding known guidance receptors. Many of these transcripts were enriched in P1 axons and are implicated in axonal growth and target recognition (Wright & Zinn, 2009). Expression levels were visualized using Z-score heatmaps generated from normalized mRNA counts across projection areas and developmental stages. This approach enabled comparison of relative mRNA abundance across regions in a standardized manner, where a Z-score of +1 reflects high expression, −1 reflects low expression, and 0 corresponds to the mean. Several guidance-related mRNAs displayed region– and stage-specific expression patterns (**Fig. 3E**). For example, Plxna4 was enriched in ribosome-bound mRNAs isolated from dorsal striatum-projecting axons at P1, while its antisense transcript, Plxna4os2, was detected in axons targeting the ventral striatum and prefrontal cortex. These findings raise the possibility of local regulatory mechanisms acting within axons. Additional guidance-related mRNAs such as Plxnc1 and DCC also showed projection-specific enrichment, consistent with previous observations (Chabrat et al., 2017; Lo et al., 2022; Reynolds et al., 2018).

Taken together, these findings suggest that ribosome-bound mRNAs in DA axons undergo dynamic changes during development, with early-stage axons enriched for transcripts involved in axon growth and guidance, and mature axons expressing mRNAs related to synaptic structure and function. This projection and stage-specific regulation reflect local translation programs that support the sequential assembly and specialization of dopaminergic circuits.

### Local translation of Plxna4 mRNA in dopaminergic axons

Among the guidance-related mRNAs identified in our transcriptomic analysis, Plxna4 emerged as a prominent candidate enriched in dorsal striatum-projecting DA axons at P1. To validate its presence in DA neurons and axons, we performed fluorescent in situ hybridization (RNAscope®) on P1 mouse brain sections. Plxna4 mRNA is detected in DA cell bodies within both the VTA and SNpc (**Fig. 4A-B**). In the striatum (**Fig. 4C-D**), a major target of DA projections, punctate Plxna4 signal is observed in both striatal neurons and TH-positive axons, indicating its presence in local cells and DA projections. To better visualize individual DA axons and confirm the axonal localization of Plxna4 mRNA, we used an *in vitro* model of E14.5 mDA explants. Fluorescent in situ hybridization (RNAscope®) was combined with immunocytochemistry against TH to label DA axons. Using this approach, we clearly detected robust Plxna4 mRNA signal within TH-positive axons (**Fig. 4E-F**), supporting its localization to the distal projections of developing mDA neurons. This distribution is consistent with a potential role for Plxna4 in axonal elongation and pathfinding during early circuit formation.

**Figure 4.**
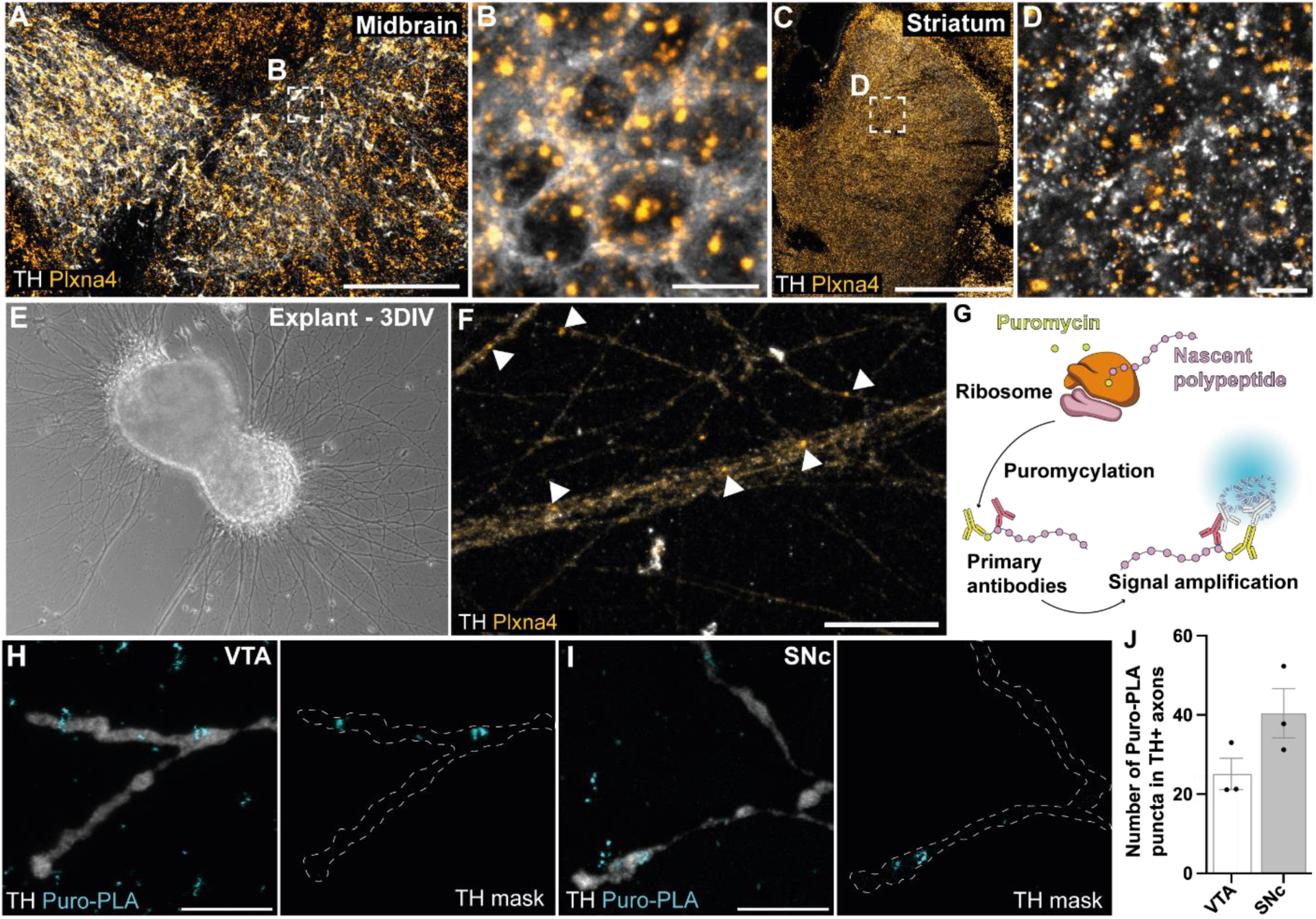
Validation of Plxna4 mRNA present in DA cell bodies and axons. A) In situ hybridization (RNAScope®) for *Plxna4* mRNA on P1 mouse midbrain section, TH immunostaining (grey) was used to reveal DA cell bodies. B) Representative high magnification of A. C) In situ hybridization (RNAScope®) for *Plxna4* mRNA on P1 mouse striatal section, TH immunostaining (grey) was used to reveal DA axons. D) Representative high magnification of C. E) Representative image of E14.5 mDA explant at 3 days *in vitro* (3DIV). F) In situ hybridization (RNAscope®) for *Plxna4* mRNA on DA axons from mDA explants, TH (grey) was used to reveal DA axons, arrows showing dots of *Plnxa4* mRNA. G) Schematic of puromycin-PLA experiments. H-I) Representative images from DA axons from VTA and SNc explants. J) Quantification of Puro-PLA puncta in DA axons reveals by TH; puncta were quantified within TH mask, and the signal was cleared outside to avoid dots from other axons.

To test whether Plxna4 mRNA is locally translated within DA axons, we employed puromycin labeling combined with proximity ligation assay (puro-PLA), a technique that detects nascent protein synthesis with subcellular resolution (De Pace et al., 2024; tom Dieck et al., 2015). E14.5 mDA explants were cultured for 3DIV, followed by a 10-minute puromycin treatment prior to fixation (**Fig. 4G**). Under these conditions, we observe Puro-PLA signal in distal axons of explants derived from both VTA and SNc regions (**Fig. 4H-J**), indicating that Plxna4 is actively translated in developing DA axons.

### Plxna4–Sema3a signaling regulates DA axonal growth in vivo and in vitro

Plxna4 is known to interact with Semaphorin 3A (Sema3a) to regulate axonal growth and guidance (Suto et al., 2005). While this interaction has been well described in sensory and cortical systems, its role in midbrain dopaminergic (mDA) neuron development remains unclear. To investigate whether Sema3a could influence mDA axon guidance, we examined publicly available in situ hybridization data from the Allen Brain Atlas. This analysis revealed strong expression of Sema3a in the pallidum and at lower levels in the striatum at E18.5 and P4 (**Fig. S4**), suggesting that Sema3a may act as a guidance cue for DA axons innervating forebrain targets during late embryonic and early postnatal stages. To investigate the function of Plxna4 during dopaminergic circuit development, we used the CRISPR-Cas9 system to knock out Plxna4 selectively in DA neurons. To confirm the efficiency of the Plxna4-targeting dgRNA, we first performed Tracking of INDELs by Decomposition (TIDE) analysis in vitro. NIH-3T3 cells were co-transfected with plasmids encoding SpCas9 and the AAV construct carrying the Plxna4-specific dgRNA, and sequencing of the targeted regions revealed indels at the expected sites, demonstrating efficient Cas9-mediated cleavage of the Plxna4 locus (**Fig. S5A, B**). We then performed *in utero* AAV injections at E12.5, a timepoint corresponding to the onset of mDA neurogenesis (Blaess & Ang, 2015), to deliver a guide RNA targeting Plxna4. We used an AAV2/Cap-B10 vector encoding mCherry and a Plxna4-specific guide RNA (AAV2/Cap-B10-mCherry-gRNA-Plxna4), injected into DAT-IRES-Cre;LSL-Cas9 embryos (Plxna4-KO). In these animals, Cas9 is selectively expressed in DAT+ neurons, allowing for cell-type-specific gene disruption. As a negative control, DAT-IRES-Cre mice lacking Cas9 were injected with the same AAV vector (control) (**Fig. 5A**) (Goertsen et al., 2022). To confirm the efficiency and specificity of the knockdown, we performed immunostaining at P8 using antibodies against TH, Plxna4, and mCherry. mCherry signal confirmed expression of the virus in TH+ neurons, and quantification revealed a ∼40% reduction in Plxna4 protein levels in Plxna4-KO mice compared to controls (**Fig. 5B-D**).

**Figure 5.**
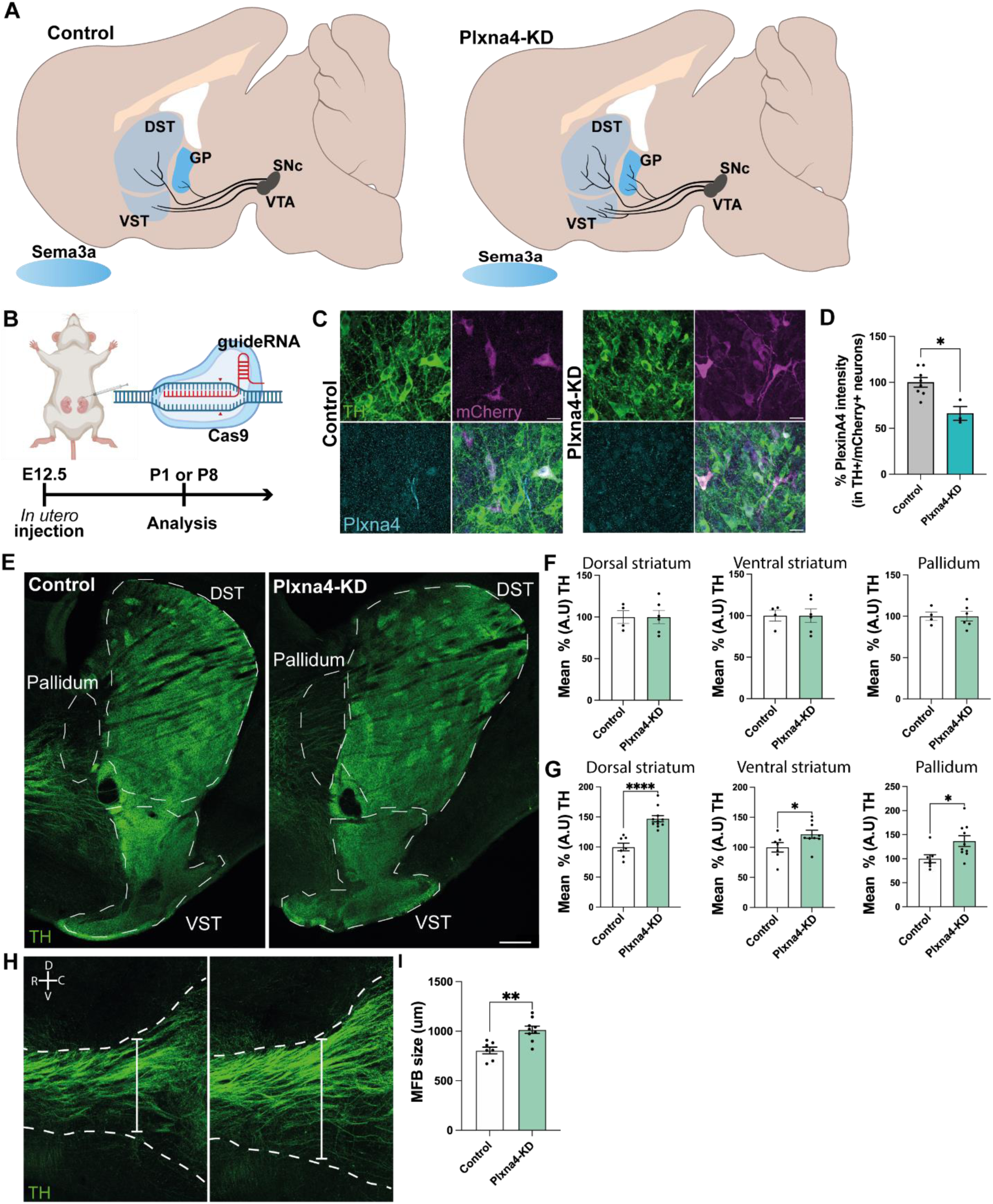
Plxna4 knock-down modulates dopaminergic axons development *in vivo*. A) Illustration of potential effect of Plxna4-KD on DA axonal growth and innervation. B) Schematic representation of the *in-utero* injection and strategy to study the impact of Plxna4-KD on DA axonal development. C) Representative high magnification images of control and Plxna4-KD animals, to visualize DA neurons TH and the viral reporter mCherry were used to assess the specific viral expression in DA neurons. D) Quantification of Plxna4 intensity in DA neurons, Plxna4 was quantified only in double positive TH/mCherry neurons. E) Representative images of dopaminergic axons revealed with a TH immunohistochemistry on P8 sagittal mouse brain sections. F) Quantification of DA axons in the dorsal striatum, ventral striatum and pallidum at one day postnatal. G) Quantification of DA axons in the dorsal striatum, ventral striatum and pallidum at eight days postnatal. H) Representative image of MFB. I) Quantification of MFB size. Statistical tests: Mann-Whitney test *p.value < 0.05 (D, G); **p.value < 0.0001 (I); ****p.value < 0.005 (G).

We next examined the consequences of Plxna4 downregulation on axonal growth and branching. At P1, TH immunostaining revealed no significant differences in axonal density in the pallidum, dorsal striatum, or ventral striatum between Plxna4-KO and control animals (**Fig. 5F**). However, by P8, a stage when mDA axons have largely completed their target innervation, TH signal intensity was significantly increased in all three regions in Plxna4-KO mice compared to controls (**Fig. 5E-G**). These findings suggest that the absence of Plxna4 results in enhanced axonal growth and arborization or possibly reduced axonal pruning during postnatal development.

To assess axon bundle organization, we examined TH+ projections in the medial forebrain bundle (MFB). Similar to phenotypes reported in mice lacking components of other semaphorin signaling pathways, such as Sema3F or Npn2 (Kolk et al., 2009), we observed axon defasciculation in Plxna4-KO animals. In Plxna4-KO mice, axon bundles appeared thicker and less compact compared to controls (**Fig. 5H-I**). Quantification revealed a significant increase in average bundle width in Plxna4-KO mice (1014 ± 37.75 µm, p < 0.0021) relative to controls (805.5 ± 33.96), consistent with reduced fasciculation in the absence of Plxna4. Together, these findings support a role for Plxna4 in restricting dopaminergic axon growth and maintaining bundle integrity during development. The increase in TH signal across forebrain targets at P8 may result from impaired Sema3a-mediated repulsion, as Sema3a is strongly expressed in the pallidum during this period. Furthermore, Plxna4 can interact with other semaphorin ligands, potentially contributing to broader effects on axonal growth and targeting.

To determine whether Sema3a acts directly on DA axons, we next performed *in vitro* assays using midbrain explant cultures from E14.5 mouse embryos. Explants from the SNpc and VTA were cultured for 3DIV and treated with Sema3a under two experimental paradigms. In a growth cone collapse assay, explants were exposed to Sema3a for 15 minutes, and the growth cone area was measured relative to untreated controls. SNpc-derived axons showed a robust collapse response, with a 56% reduction in growth cone area following treatment. In contrast, VTA-derived axons were unaffected (**Fig. 6A-C**). To assess longer-term effects on axonal branching, we performed a Sholl analysis after treating explants with a lower concentration of Sema3a for 24 hours. Sema3a significantly reduced axonal arborization in SNpc-derived explants, while no effect was observed in VTA-derived cultures (**Fig. 6D-G**).

**Figure 6.**
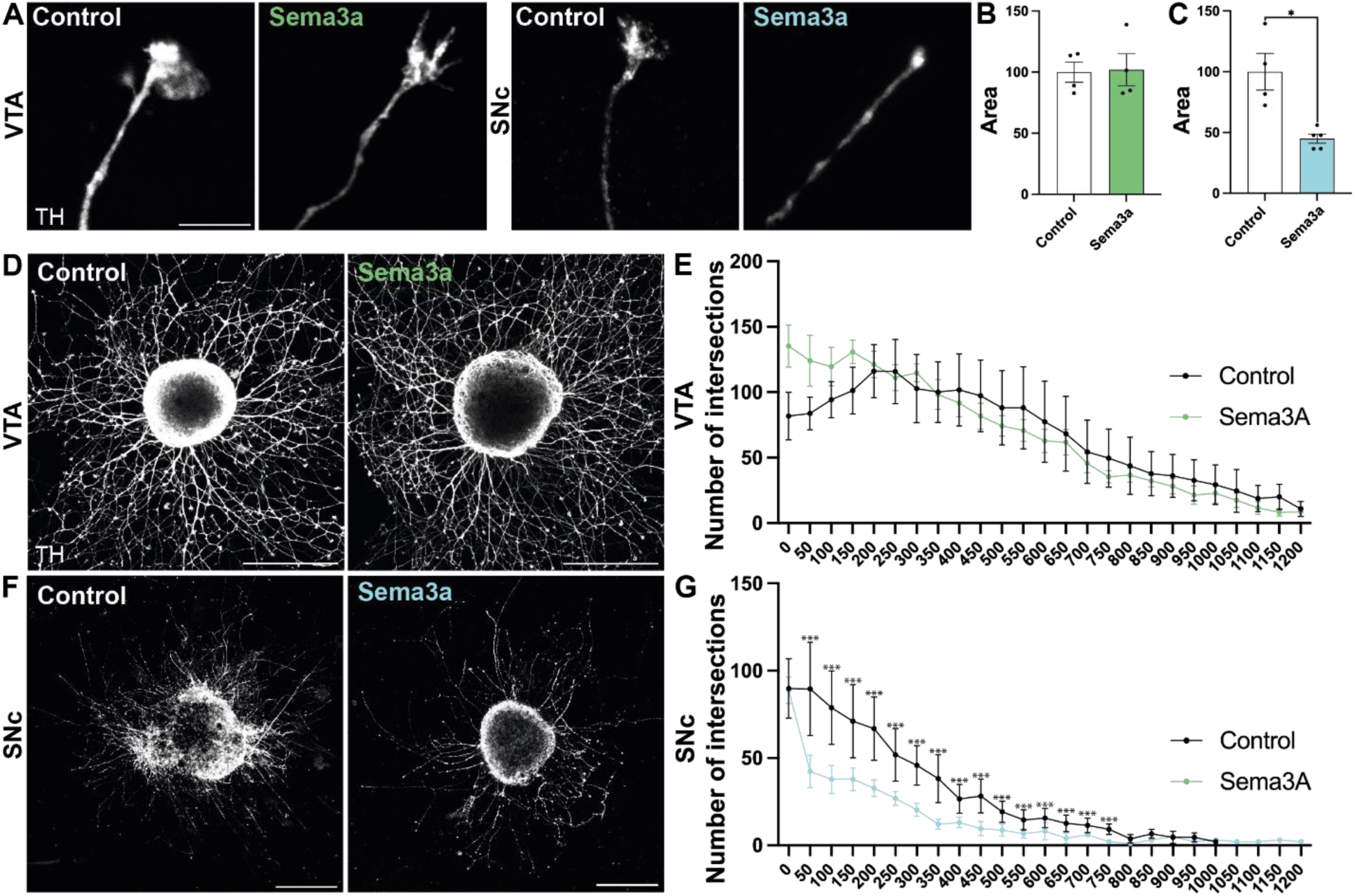
Sema3a influences Plxna4 expression in growth cones and reduces axonal outgrowth *in vitro*. A) Representative images of DA growth cones in SNc and VTA explants with (control) or without Sema3a treatment from 15 minutes, TH was used to mark DA axons. B-C) Quantification of growth cones area of DA axons from VTA explants (B) and SNc explants (C), GC area was normalized on control growth cones. D) Representative images of VTA explants with or without Sema3a treatment for 24hours, TH was used to label DA axons. E) Quantification of axon arborization from VTA explants with or without Sema3a exposure. F) Representative images of SNc explants with or without Sema3a treatment for 24hours, TH was used to reveal DA axons. G) Quantification of axon arborization from VTA explants with or without Sema3a exposure. Statistical tests: Mann-Whitney test *p.value < 0.05 (C); Two-way ANOVA with multiple comparisons ***p.value < 0.005

These in vitro findings confirm that Sema3a can directly influence mDA axon morphology in a region-specific manner and support a functional interaction with Plxna4. Taken together with our *in vivo* results, these data establish Plxna4–Sema3a signaling as a key modulator of axonal growth and fasciculation in the developing dopaminergic system.

## DISCUSSION

Our study reveals that local mRNA translation is an integral mechanism shaping DA circuit development. Using a RiboTag strategy combined with DAT-IRES-Cre, we confirmed that ribosomes are present in DA axons, providing direct evidence for local protein synthesis in this system. As in other neuronal populations, ribosome levels were considerably lower in axons compared to somata (Fusco et al., 2021; Koppers et al., 2024; Nagano & Araki, 2021), which may explain why earlier work reported negative results (Hobson et al., 2022). By applying more sensitive methods, including ribosome immunoprecipitation and proximity ligation assays, we could detect HA-tagged ribosomes in DA axons. We further demonstrated the presence and active translation of axonal mRNAs using fluorescent in situ hybridization and puromycin labeling. Together, these data provide compelling evidence that DA axons contain functional ribosomes and engage in local protein synthesis during development.

By profiling axonal translatomes at multiple developmental stages and across projection targets, we uncovered a dynamic and compartment-specific regulation of translation in DA neurons. Early postnatal axons were enriched in mRNAs encoding proteins for cytoskeletal remodeling, axonogenesis, and guidance, consistent with their ongoing growth and pathfinding. In contrast, mature axons exhibited enrichment for transcripts associated with synaptic function, neurotransmitter release, and plasticity. These temporal shifts mirror the evolving functional demands of axons, first supporting elongation and targeting, then stabilizing connections and enabling synaptic modulation. The fact that these developmental changes were far more pronounced in axons than in somata underscores the importance of spatially localized translation in coordinating circuit assembly.

A second key finding is that axonal translatomes are highly projection-specific. Distinct sets of ribosome-bound mRNAs were identified in axons projecting to the dorsal striatum, ventral striatum, and prefrontal cortex, with only limited overlap across targets. This compartmentalization suggests that axonal translation is tailored to the requirements of each projection, potentially contributing to the topographic organization of DA circuits. Such compartment-specific translation may represent a mechanism by which diverse DA subtypes, with shared somatic transcriptomes, acquire distinct connectivity patterns.

Among the guidance-related transcripts selectively enriched in dorsal striatum-projecting axons, Plxna4 emerged as a compelling candidate. We confirmed its localization in DA axons using *in situ* hybridization in both brain tissue and explant cultures and demonstrated its local translation with puromycin-PLA. These findings extend earlier work showing axonal enrichment of guidance transcripts in other neuronal systems (Koppers et al., 2019; Lu et al., 2024; Shigeoka et al., 2016; Zivraj et al., 2010) and provide direct evidence that DA axons rely on local synthesis of guidance receptors during development.

Functional experiments revealed that Plxna4–Sema3a signaling is a critical regulator of DA axon morphology. In vitro, Sema3a exposure induced collapse and reduced arborization in SNc-derived, but not VTA-derived, axons, highlighting region-specific sensitivity to this cue. In vivo, partial Plxna4 knockdown increased TH+ axonal density in both striatum and pallidum and led to defasciculation in the medial forebrain bundle. Together, these results position Plxna4–Sema3a signaling as a central mechanism that constrains DA axon growth, maintains bundle integrity, and ensures correct target innervation. The region-specific effects observed in vitro further suggest that Plxna4-dependent responses are tuned according to DA neuron subtype, consistent with emerging evidence of molecular diversity within this system (Azcorra et al., 2023; Gaertner et al., 2022).

While our findings advance the understanding of DA axon development, certain technical considerations remain. The RiboTag approach captures only ribosome-associated mRNAs bound to tagged subunits and may underrepresent transcripts undergoing non-canonical or stalled translation (Fusco et al., 2021). Moreover, our use of heterozygous RiboTag animals limited HA-ribosome abundance, and homozygous lines could improve yield and reduce RNA-seq variability.

Taking together, our results demonstrate that local mRNA translation is not a peripheral process but a central mechanism guiding DA circuit assembly. The identification of projection-specific translatomes and the functional role of Plxna4–Sema3a signaling highlight how spatially restricted protein synthesis contributes to axonal growth, branching, and targeting. These findings extend the concept of axonal translation from sensory and cortical neurons to the dopaminergic system and establish a framework for understanding how local translation programs encode projection-specific identities.

Beyond development, these results hold relevance for disease and repair. Aberrant DA axonal connectivity has been implicated in neurodevelopmental disorders such as autism and schizophrenia, while progressive degeneration of nigrostriatal projections is the hallmark of Parkinson’s disease. Our data suggest that strategies designed to modulate local translation, or to harness endogenous guidance pathways, may improve the integration of grafted DA neurons into host circuitry. For example, guiding transplanted neurons to re-establish precise nigrostriatal connectivity may depend not only on extrinsic cues but also on the intrinsic ability of axons to locally synthesize guidance receptors such as Plxna4.

In summary, our study demonstrates that dopaminergic axons harbor ribosomes and actively translate mRNAs in a projection– and stage-specific manner. We show that the axonal translatome undergoes a developmental shift, from guidance– and growth-related programs in early postnatal stages to synaptic– and plasticity-related functions in adulthood. Among guidance-related transcripts, Plxna4 stands out as locally translated in DA axons, and our functional data indicate that Plxna4–Sema3a signaling contributes to the regulation of axon growth, branching, and fasciculation. These results highlight local translation as one of the molecular layers shaping DA circuit formation. Beyond Plxna4, we also identified other guidance-related mRNAs, including members of the Semaphorin, Ephrin, and Slit families, whose local translation may contribute to the precise establishment of DA connectivity and represent important candidates for future investigation. Future studies will be needed to dissect the mechanisms governing axonal mRNA localization and translation, and to determine how these processes influence circuit stability in the adult brain and in disease contexts such as Parkinson’s disease.

## RESOURCE AVAILABILITY

Lead contact: Further information and requests for resources and reagents should be directed to and will be fulfilled by the lead contact, Martin Lévesque.

Materials availability: This study did not generate new unique reagents. Plasmids and viral constructs used in this study are available from the lead contact upon reasonable request and completion of a materials transfer agreement (MTA).

RNA sequencing data generated in this study have been deposited in ArrayExpress under accession number E-MTAB-16070. The dataset is accessible at: https://www.ebi.ac.uk/biostudies/arrayexpress/studies/E-MTAB-16070.

This paper does not report original code.

Any additional information required to reanalyze the data reported in this paper is available from the lead contact upon request.

## ACKNOWLEDGEMENTS

The authors thank the CERVO Centre Molecular Tool Platform (https://neurophotonics.ca/fr/pom) for the production of viral vectors. This work was supported by research grants from the Natural Sciences and Engineering Research Council of Canada (NSERC, RGPIN-2024-05363 to M.L. and RGPIN-2020-06376 and DGECR-2020-00060 to C.F.S) and the Canadian Institutes of Health Research (CIHR, 451548 to M.L.). M.L. is the recipient of a senior career award from the Fonds de Recherche du Québec–Santé (FRQS) in partnership with Parkinson Québec (34974). E.M was funded by the UK DRI RRZA-175 grant.

## AUTHOR CONTRIBUTIONS

C.G., and M.L. designed the experiments of the study. C.G. performed all in vitro and in vivo experiments, prepared the manuscript with edits by M.L. B.F and S.H. performed bioinformatic analysis. E.M co-supervised the work and performed mRNA sequencing. P.G and C.F.S polyribosomes extraction. All authors approved the final version.

## SUPPLEMENTAL INFORMATION

Supplemental information includes Document S1, which contains Figures S1–S5 and their corresponding figure legends.

## METHODS

### Animals

All animal experiments were performed in accordance with the Canadian Guide for the Care and Use of Laboratory Animals and were approved by the Laval University Animal Protection Committee. *Dat^Cre/+^*(Zhuang et al., 2005) and *Pitx3^Gfp/Gfp^* (Zhao et al., 2004) mice were genotyped as previously described. Dat-IRES-Cre × RiboTag animals were generated by intercrossing Dat-IRES-Cre^het^ male (Jaxson 006660) and RiboTag^ho^ female (Jaxson 011029) animals. Transgenic mice were identified by PCR using forward primer (5’-ATCCGAAAAGAAAACGTTGA-3’) and reverse primer (5’-ATCCAGGTTACGGATATAGT-3’) targeting the Cre sequence.

### Tissue analysis

Brains from P1 mice were fixed in 4% paraformaldehyde (PFA) in 1× PBS at 4 °C, followed by cryoprotection in 30% sucrose in PBS and freezing on dry ice. For mice older than P1, intracardiac perfusion with 4% PFA in PBS was performed prior to cryoprotection. Brains were sectioned at 50 μm on a cryostat, washed in PBS, and blocked with 1% normal donkey serum (NDS) and 0.2% Triton X-100 for one hour. Primary antibodies used were: rabbit anti-TH (Pel-Freez, 1:1000), sheep anti-TH (Millipore, 1:1000), rabbit anti-HA (Sigma, 1:500), mouse anti-HA (Covance, 1:1000), rabbit anti-Plxna4 (Abcam, 1:500), and chicken anti-mCherry (Aves Labs, 1:200). Secondary antibodies (donkey Alexa Fluor 488, 555, 594, and 647) were used at 1:400.

For fluorescent in situ hybridization (RNAscope®) on tissue, P1 brains were fixed in 4% PFA in DEPC-treated PBS at 4 °C, cryoprotected in 30% sucrose, and frozen on dry ice. Brains were sectioned at 30 μm and collected on Superfrost Plus slides (Fisher Scientific). RNAscope probes for mouse Plxna4 (515491-C2) were obtained from Advanced Cell Diagnostics (Newark, CA, USA). In situ hybridization was performed according to the manufacturer’s protocol.

### mRNA sequencing

*Sample collection*. Brain tissue from the midbrain, dorsal–ventral striatum, and prefrontal cortex (PFC) was freshly dissected and flash-frozen at –80 °C. Tissue was collected at two developmental stages: P1 and adult (>P60). Three animals per stage were used, yielding twelve samples per stage.

*RNA extraction, immunoprecipitation, and purification*. RNA was extracted, immunoprecipitated, and purified as described previously (Shigeoka et al., 2016) with minor modifications. Briefly, frozen tissue was homogenized in ice-cold CHX lysis buffer (20 mM HEPES-KOH, 5 mM MgCl₂, 150 mM KCl, 1 mM DTT, 1% NP-40, 200 U/mL RNase inhibitor [NEB M0314], 100 μg/mL cycloheximide, and protease inhibitor cocktail). Lysates were centrifuged at 16,000 × g for 10 min at 4 °C. The supernatant containing tagged ribosome–mRNA complexes was incubated with magnetic Protein A/G beads (Thermo Scientific Pierce) and rotated for 1 h at 4 °C. Precleared lysates were incubated overnight with 2.5 μL anti-HA antibody (Abcam, ab9110), then pulled down with Dynabeads. Beads were washed, and RNA was eluted with CHX lysis buffer supplemented with RTL buffer (Qiagen). RNA was purified using the RNeasy Mini Kit (Qiagen).

*RNA sequencing*. RNA-seq libraries were generated from immunoprecipitated fractions using the Ovation RNA-Seq System V2 (Tecan) according to the manufacturer’s protocol, at the laboratory of E. Metzakopian (UK Dementia Research Institute, University of Cambridge).

*Bioinformatic analysis*. Read quality was assessed using FastQC. Reads were trimmed with Trim Galore (v0.6.5) and pseudo-aligned with Kallisto (v0.46.1)(Bray et al., 2016) to the ENSEMBL GRCm39 mouse transcriptome (mm39, release M36) (Mudge et al., 2025). Between 80–94% of reads mapped successfully (Supplementary Table S1). Transcript-level abundance was imported and summarized at the gene level with tximport (v1.43.0)(Soneson et al., 2015). Genes with fewer than 1 count in at least 2 replicates across all groups were removed. DESeq2 (v1.46.0) was used for normalization and differential expression analysis. Genes with ≥1.5-fold change and adjusted p-value <0.05 were considered differentially expressed. Gene Ontology enrichment was performed with clusterProfiler (v4.14.6)(Yu et al., 2012).

### Polyribosomes extraction

Frozen midbrain and striatum tissues (pooled from 4 transgenic animals per region at P1) were homogenized in ice-cold 2x polyribosome lysis buffer (20 mM Tris-HCl pH 7.4, 5 mM MgCl₂, 100 mM KCl, 1% NP-40, 1 mM DTT, 20 U/mL SUPERase-In, protease inhibitors [EDTA-free], PhosSTOP, and 100 μg/mL cycloheximide) using a Dounce homogenizer. Homogenates were centrifuged at 1,200 × g for 5 min at 4 °C. Supernatants were loaded onto 10–50% (w/v) sucrose gradient and ultracentrifuged (222 000 × g, 2 h 45 min, 4 °C, SW41 Ti rotor, Beckman Coulter). Gradients were fractionated using a BR-188 system (Brandel) with UV monitoring at 254 nm. Fractions were ethanol-precipitated overnight at –20 °C, pelleted, resuspended in Laemmli buffer, and boiled for 5 min at 95 °C (Sevigny et al., 2020).

### Western blotting

Proteins were resolved by SDS-PAGE (10%) and transferred to 0.45 μm nitrocellulose membranes. Membranes were blocked in 5% milk or BSA in TBS-T (10 mM Tris-HCl pH 7.6, 150 mM NaCl, 0.1% Tween-20) for 1 h at room temperature, incubated overnight with primary antibodies at 4 °C, and with secondary antibodies for 1 h at room temperature. Blots were imaged using the Odyssey Imaging System (Li-COR Biosciences).

### *In vitro* mDA explant cultures

*mDA explant preparation*. Midbrain explants were dissected at E14.5 from Pitx3 Gfp/+ embryos in ice-cold L-15 medium with 5% FBS. Explants were cultured for 3 DIV in ibidi 8-well chambers coated with laminin (20 μg/mL, 2D cultures for RNAscope® and Sholl analysis) or embedded in Matrigel (3D cultures for collapse assay and puromycin-PLA). Media consisted of neurobasal medium supplemented with B27, GlutaMAX, sodium pyruvate, Pen/Strep, 0.4% methylcellulose, and 5% FBS.

*Fluorescent in situ hybridization (RNAscope®).* Explants were fixed after 3 DIV in 4% PFA (DEPC-treated) with 4% sucrose in PBS for 30 min at 4 °C. RNAscope probes for mouse *Plxna4* were used according to the manufacturer’s instructions.

*Collapse assay.* Explants were treated with Sema3a (250 ng/mL) or PBS for 15 min in serum-containing medium, then fixed as above. TH was immunostained with sheep anti-TH (Pel-Freez, 1:1000) and Alexa Fluor 488 anti-sheep secondary (1:400). Growth cone area was quantified in ImageJ.

*Sholl analysis.* Explants from VTA and SNc were treated with Sema3a (100 ng/mL) or PBS for 24 h in serum-free medium, then fixed and TH-immunostained as above. Axonal arborization was quantified with the ImageJ Neuroanatomy plugin. Radii were defined at 10 μm intervals, and intersections were averaged across samples and plotted every 50 μm.

*Puromycin-PLA.* Explants were incubated with puromycin (2 μM, 10 min), washed, and fixed in 4% PFA with sucrose. Controls were fixed without puromycin. Blocking, primary antibody incubation (anti-Plxna4 and anti-puromycin), PLA probe hybridization, ligation, amplification, and detection were performed with the Duolink® kit (Sigma-Aldrich) according to the manufacturer’s instructions.

### Plxna4 dgRNA validation (TIDE)

Tracking of INDELS by Decomposition (TIDE) was used to validate gRNA efficiency (Brinkman & van Steensel, 2019). NIH-3T3 (ATCC, CRL-1658) were cultured in Dulbecco’s modified Eagle’s Medium (DMEM, Sigma) supplemented with 10% fetal bovine serum and 1% penicillin-streptomycin (Thermo Fisher Scientific). The cells were co transfected with Lipofectamine LTX (Invitrogen, 15338-030) with plasmid encoding Streptococcus pyogenes Cas9 (SpCas9) (Addgene, plasmid #62988) and with plasmid CAG-DIO-mcherry-U6 dgRNA-Plxna4 (gRNA1: GCCACACTCGAAATCCGGGT; gRNA2: GTTGCCACTGGTGTAAATAC) or with the control plasmid CAG-DIO-mCherry. Transfected cells were selected with 2.0 µg/µl puromycin. gDNA was extracted using Monarch® Spin gDNA Extraction Kit (New England Biolabs, T3010). The exons 12 and 9 of Plxna4 were amplified by PCR using Q5® High-Fidelity DNA Polymerase (New England Biolabs, M0491). The exon 12 of Plxna4 was amplified using the primers (F)CTGAAGTGGGTTACAGGTGGGACAG and (R)CCTCGCTCGGTTCCATCCACTTTTG, whereas the exon 9 was amplified with the primers (F)TATGCACCACACCTGAAGAGCCAAG and (R)GATCAACAATGGCTGGAGAGGGAGG.

PCR program: initial denaturation at 95°C for 4 min, denaturation at 94°C for 30 sec, annealing at 72°C for 45 sec, elongation at 72°C for 1 min, with 30 cycles. The amplicons were sequenced by SANGER at the sequencing platform from CHU of Laval University.

### Statistical analysis

Data were analyzed with GraphPad Prism 10.4.9 (GraphPad Software, La Jolla, CA, USA). Collapse assay, TH intensity, and medial forebrain bundle (MFB) size were analyzed using two-tailed Mann–Whitney tests. Sholl analysis was analyzed by two-way ANOVA with multiple comparisons. A significance threshold of p < 0.05 was used.

### Microscopy

Images were acquired using a Zeiss LSM700 confocal microscope and processed with ImageJ. Experiments included fluorescent in situ hybridization (RNAscope®), collapse assays, puromycin-PLA, and immunofluorescence.

